# A novel miniaturized filamentous phagemid as a gene delivery vehicle to target mammalian cells

**DOI:** 10.1101/2024.10.30.621196

**Authors:** Shirley Wong, Salma Jimenez, Deborah Pushparajah, Rohini Prakash, Roderick Slavcev

**Affiliations:** School of Pharmacy, University of Waterloo, Canada; Centre for Eye and Vision Research (CEVR), 17W, Hong Kong Science Park, Hong Kong

**Keywords:** filamentous bacteriophage M13, phagemid, gene transfer, non-viral gene delivery, DNA minivector

## Abstract

The filamentous phage M13 is a single-stranded DNA phage with several attractive characteristics for gene delivery, including a capsid amenable to the display of foreign peptides, and a simple well-characterized genome that is easy to genetically modify. Previously, we constructed a DNA minivector based on M13 (a miniphagemid), which minimized the inflammatory bacterial and phage DNA content in the vector. In general, DNA minivectors, devoid of their prokaryotic components have shown improved gene transfer and safety. We examined the miniphagemid’s capacity for *in vitro* transgene delivery to target cells through phage display of epidermal growth factor to target its cognate receptor. The absence of the prokaryotic backbone and smaller vector size conferred by the miniphagemid was associated with improved transgene expression for purified single-stranded phagemid DNA and phagemid virion particles. We further engineered this system to enhance packaging of DNA minivectors by deletion of the packaging signal within the helper plasmid, used to produce miniphagemids, and observed improved phage-mediated gene expression in mammalian cells. Overall, here, we present a set of novel transgene delivery vectors that combine cell-targeting ligand display and vector minimization. This platform showcases the flexibility of M13 as a gene delivery tool with immense therapeutic potential.

## Introduction

Viruses, such as the filamentous bacteriophage (phage) M13, exist as inert proteinaceous particles outside their host bacterium, *Escherichia coli (E.coli)*. As they do not possess intrinsic tropism for mammalian cells, they can function as simplistic carriers of genetic material for gene transfer applications. However, phages also have no intrinsic means by which to transduce mammalian cells. Indeed, the most common gene transfer vectors are modified viruses of mammalian and human origin^1,2^ and exhibit natural tropism for human cells. However, some safety and efficacy concerns limit their usage, specifically, high immunogenicity, risk of insertional mutagenesis, and the possibility of recombination into replication-competent viruses.^3–6^ In contrast, non-viral methods of gene transfer incorporate other means of bypassing cellular barriers for gene transfer, such as with chemical transfection reagents that mediate transport across cell membranes.^7,8^ However, these transfection reagents cannot target specific tissues; functionalization requires additional covalent linkage which may not be amenable to scale-up.^7^

As phages do not innately enter, nor replicate within mammalian cells, they may be considered as “non-viral” gene transfer vectors. To overcome their lack of tropism, display of a cell-targeting ligand on a phage capsid can facilitate cellular uptake by exploiting receptor-mediated endocytosis.^9–11^ Genetic incorporation of a sequence for phage display easily facilitates decoration of the phage capsid with any ligand of choice. The filamentous phage M13 is an excellent model for the display of cell-specific ligands. The phage’s simplistic genome encodes all proteins necessary for replication, progeny assembly, and extrusion, which are controlled by signalling structures within the phage replicative origin (f1 *ori*) in the genome. The f1 *ori* contains all sequences necessary and sufficient to direct phage-mediated replication of a DNA molecule. Therefore, any conventional plasmid that also contains an f1 *ori,* in addition to its plasmid *ori,* can be replicated by filamentous phage machinery independent of its plasmid origin^12,13^ and assembled into virion particles.

The prokaryotic backbone is necessary for amplification and maintenance in a bacterial host, however, it is rich in unmethylated cytosine-guanine dinucleotide (CpG) motifs that are known to inhibit transgene expression in mammalian cells.^14^ Indeed, CpG-mediated immunostimulation constitutes a major part of the immune response against M13 administered in mammals.^15,16^ The prokaryotic backbone of a typical plasmid used for gene delivery furthermore contains an antibiotic resistance marker that can disseminate antibiotic resistance into the environment.^17^ Ultimately, this backbone does not contribute at all to transfection and is thus unnecessary bulk. We and others have documented efforts in producing precursor vectors for M13-mediated production of DNA minivectors (Supplementary Figure 1) deficient in bacterial or phage genetic sequences.^18,19^ Here, we evaluated if filamentous M13 miniphagemid particles can improve phage-mediated delivery of a mammalian transgene cassette, *cmv-luc,* based on the display of a cell-specific ligand for internalization and the absence of the prokaryotic backbone. We also further engineered vectors for M13-mediated production to improve DNA minivector packaging by limiting the packaging of full precursor DNA and helper plasmid, in order to investigate if this could consequently enhance gene delivery and expression in mammalian cells.

## Materials and Methods

### Strains and vectors

*E. coli* K-12 JM109 was used in the generation of all phage and plasmid constructs. All bacterial and mammalian cell lines are listed in Supplemental Table 1, plasmids in Supplemental Table 2, and phages in Supplemental Table 3. All mammalian cell lines were maintained in tissue culture plates (Thermo Scientific) at 37 °C in a humidified atmosphere with 10% CO_2_ and cultured in Dulbecco’s Modified Eagle’s Medium (DMEM) (Thermo Scientific) supplemented with 10% heat-inactivated fetal bovine serum (FBS) and 1% penicillin/streptomycin. Bacterial strains were cultured in Luria–Bertani (LB) liquid medium, supplemented with the relevant antibiotic as required. The list in Supplemental Table 2 includes the plasmids that were tested for packaging within the produced full and miniphagemids, in addition to helper plasmids which were constructed as outlined in the subsequent section. The list in Supplemental Table 3 outlines the full and miniphagemids that were produced in this study. Phages were purified through precipitation with polyethylene glycol (PEG) and stored in Tris-NaCl (TN) buffer as previously described.^18^

### Construction of M13KO7 derivatives with pIII::EGF fusion display

M13KE has endonuclease target sites in gIII (KpnI-EagI) that simplify N-terminal peptide fusions^20^, while M13KO7 contains the plasmid p15a *ori* for phage-independent amplification and a KanR marker for antibiotic selection.^21^ The *gIII* from M13KE was inserted into the helper phage M13KO7 using Gibson assembly to generate the helper phage M13SW7, which can easily take on N-terminal pIII fusions while retaining the p15a *ori* to simplify selection and amplification. Primers are summarized in Supplemental Table 4. Next, the peptide EGF was inserted as a pIII N-terminal fusion with a GGGS (Gly-Gly-Gly-Ser) linker^22^ between the KpnI-EagI sites of M13SW7. Insertion was verified in the final construct (M13SW7-EGF) through PCR of the phage lysate. Peptide display of EGF was verified through dot blot and ELISA using anti-EGF antibodies (Thermo Fisher Scientific). An additional self-packaging deficient variant of M13KO7 (M13SW8) was engineered with a deleted packaging signal (PS) and pIII:EGF fusion display, as previously described.^18^ Display of the EGF peptide was completed in the same manner as M13SW7-EGF, to generate M13SW8-EGF.

### Double-stranded DNA and single-stranded miniphagemid DNA purification

These steps were completed as previously outlined.^18^

### Assessment of M13SW8 packaging efficiency

Colony assay spot plating of M13SW8-mini-(luc) and M13SW8-mini_egf_-(luc) was conducted. This was conducted to assess for packaging contamination of helper plasmids (M13SW8 and M13SW8-EGF) and full precursor plasmid (pM13ori2.cmvluc) by evaluating the ability of the produced phagemids to confer antibiotic resistance to susceptible cells. Considering pM13ori2.cmvluc contains an ampicillin resistance gene and the helper phage plasmids (M13SW8 and M13SW8-EGF) contain a kanamycin resistance gene, any phagemid growth on ampicillin, kanamycin and ampicillin + kanamycin LB plates demonstrate packaging contamination, as miniphagemids should only package the gene cassette of interest. To conduct the colony assay spot plates, 200 uL of *E.coli* JM109 cells were added to 3 mL top agar supplemented with 5mM MgSO_4_ and poured onto pre-warmed LB plates with relevant antibiotics (ampicillin, kanamycin, and ampicillin + kanamycin). 10 uL of phagemids, diluted in TN buffer from 10^-2^ to 10^-8^ were plated. The plates were incubated overnight at 37 °C. Percentage of efficiency of plating was then calculated by the following equation: (contaminated phagemid (helper phage + full precursor) titre / total phagemid titre) x 100.

### Assessment of cell viability after exposure to phage particles

HeLa cells were seeded in 96-well plates (Thermo Fisher Scientific) at 5 × 10^4^ cells/mL. The following day, EGF-displaying (M13SW7-EGF) and non-displaying (M13KO7) phage were transfected into each well at concentrations between 5 × 10^7^ virions/mL. Cell viability was assayed using 3-(4,5-dimethylthiazol-2-yl)-2,5-diphenyltetrazolium bromide (MTT) after 24 and 96 h by measuring A_492_ on a Varioskan LUX multimode plate reader (Thermo Fisher Scientific). Cell viability was reported as a percentage of the difference between the absorbances of each sample (A_sample_) and negative control (A_negative_) relative to the absorbance of the untreated control, A_NTC_: (A_sample_ – A_negative_) / A_NTC_.

### Localization of transfected phage particles

PEG-precipitated M13SW7-EGF (EGF^+^) and M13KO7 (EGF^−^) were labeled with Alexa Fluor 488 (Thermo Fisher Scientific). HeLa cells were seeded in 24-well plates at 5 × 10^4^ cells/mL. The following day, labeled phage particles and mini-(*gfp*) were transfected into cells at 5 × 10^7^ virions/mL. Wells were imaged 1h, 6 h, 24 h, 48 h, 72 h, and 96 h after transfection. To image, cells were first fixed with 4% para-formaldehyde and permeabilized with 0.1% Triton-X. Nuclei were stained with 4′,6-diamidino-2-phenylindole (DAPI), while actin was stained with rhodamine phalloidin. Fixed cells were imaged on the EVOS FL Auto Imaging System (Thermo Fisher Scientific) at 40X magnification using the DAPI (nuclei), red fluorescent protein (RFP) (actin), and GFP (phage or expressed GFP) channels.

### Transfection of phage in EGFR^+^ cell lines

Cell lines were seeded at the following cell densities in 24-well plates: 5 × 10^4^ cells/mL (HeLa), 1 × 10^5^ cells/mL (HEK293T, MRC-5, HT-29). Phage particles were added to a final concentration of 5 × 10^7^ virions/mL of phagemid. Helper phage alone was transfected as a negative control. To assess the influence of a cationic polymer transfection carrier, phage particles were also complexed with 2 μL of TurboFect. Transfection was quantified through luminescence (Luciferase Reporter Assay System; Promega) after 96 h. Luminescence was normalized against whole protein content, which was estimated via a bicinchoninic acid (BCA) assay (Thermo Fisher Scientific). The efficiency of gene transfer was reported as luminescence per 100 μg of whole protein content (RLU/100 μg).

### Phosphorylation of EGFR in HeLa

Activation of EGFR, as assessed by phosphorylation of the tyrosine residue at 1173, was visualized via Western blot. HeLa cells were treated with EGF^−^ phage (M13KO7), recombinant EGF (Thermo Fisher Scientific), or EGF^+^ phage (M13SW7-EGF) for 5 min prior to cell lysis. This was repeated in cells pre-treated with EGFR inhibitor gefitinib (Cell Signaling Technologies) 2 h prior to addition of EGF or M13SW7-EGF. The Western blot was performed using anti-EGF antibodies as outlined above, including the use of β-actin as a loading control. Lysates were probed for phosphorylated EGFR using a phospho-specific EGFR rabbit mAb specifically against Tyr1173 (53A5, Cell Signaling Technology).

### Statistical analysis

Values are reported as means of *n* independent experiments with uncertainty reported as the standard deviation (SD). Statistical hypothesis tests were evaluated using one-way ANOVA, followed by the Tukey range test for multiple comparisons. Values of *p* < 0.05 were considered statistically significant. The phagemid fraction was determined as the concentration of target phagemid divided by the total virion concentration, expressed as percentages. As compositional data^24^, they have a fixed constant sum constraint (100%). In order not to violate this constraint, the data were transformed using an isometric log ratio transformation before performing statistical analyses and transformed back to percentages for reporting.

## Results

### Display of a cell-specific ligand

A helper phage derivative of M13KO7 and M13KE (M13SW7) was constructed by replacing the *gIII* gene in M13KO7 with the display-accommodating *gIII* from M13KE. The unique endonuclease target sites (KpnI and EagI) in M13KE-derived *gIII* easily facilitate in-frame N-terminal fusions of a display peptide. Genetic insertion of *egf* with a Gly-Gly-Gly-Ser linker (*egf::GGGS*) between KpnI and EagI was verified by PCR (Figure 1A), while peptide display of the EGF peptide on the virion was verified by dot blot and ELISA (Figure 1B,C). M13SW7-EGF titres, as measured by plaque assay (Supplemental Table 5), were also comparable to those of its predecessors, M13SW7 and M13KO7, as well as wild-type M13, indicating that the EGF fusion was well-tolerated by the phage and did not impede infectivity. No significant difference was observed in packaging efficiency by phage displaying or not displaying EGF (Figure 2). Instead, packaging efficiency was largely determined by the phagemid. Miniaturized phagemids were more preferentially packaged over their full phagemid counterparts.

**Figure 1.**
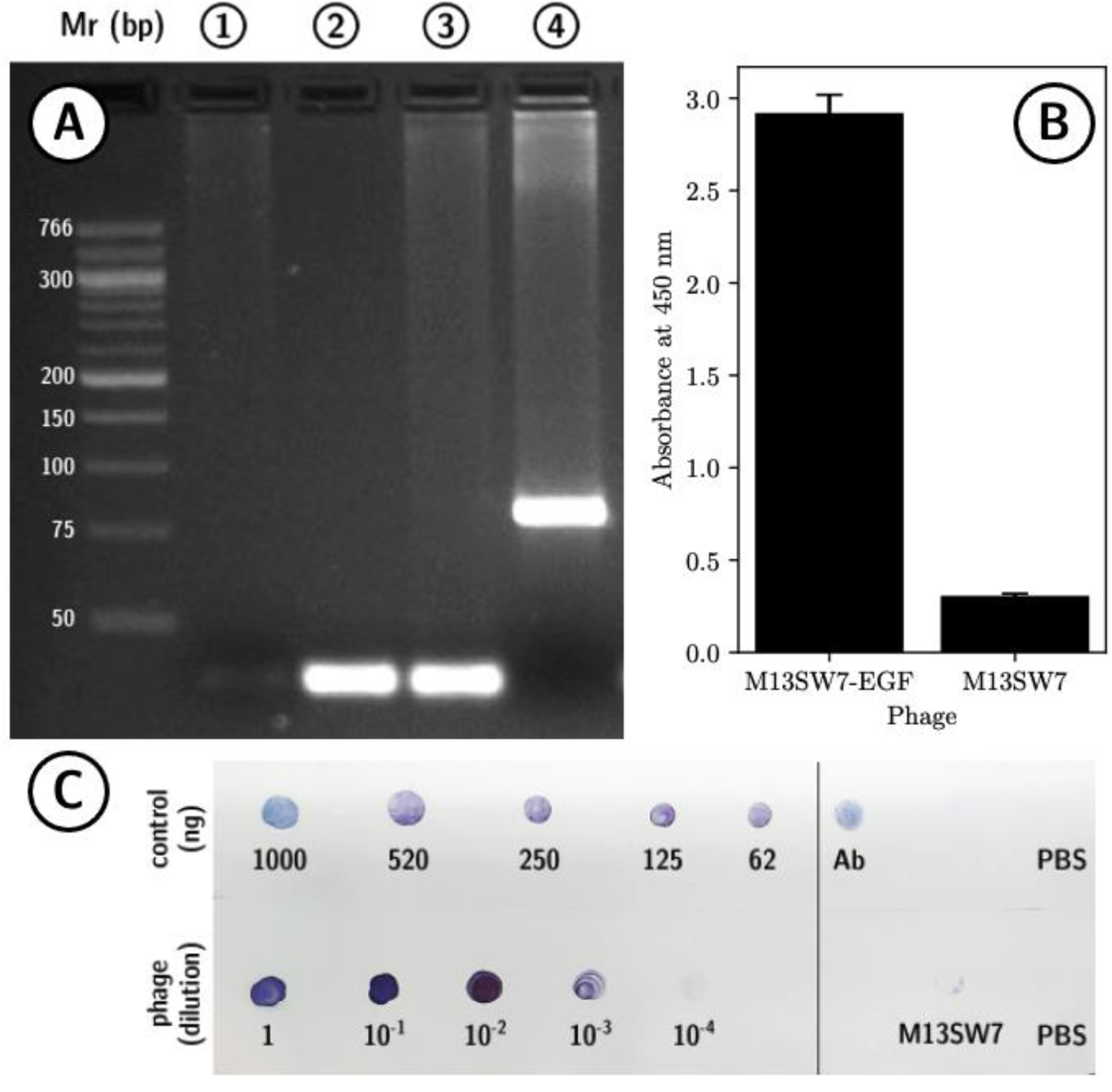
EGF display is confirmed on recombinant helper phage. A) The KpnI-EagI region in *gIII* was amplified by PCR and visualized via AGE. From left to right: M13KO7 (negative control, no KpnI-EagI region), M13KE and M13SW7 (no EGF; 54 bp KpnI-EagI region), 4) M13SW7-EGF (228 bp), Mr: Low Molecular Weight Ladder (New England BioLabs). B) ELISA of the putative EGF^+^ helper M13SW7-EGF is compared to the EGF^−^ precursor, M13KO7. Error bars represent SD, *n*=3. C) A dot blot comparing recombinant EGF (control, top) to M13SW7-EGF (bottom); controls: antibody spotted directly on the membrane (positive), EGF^−^ phage (negative) and phosphate-buffered saline (negative).

**Figure 2.**
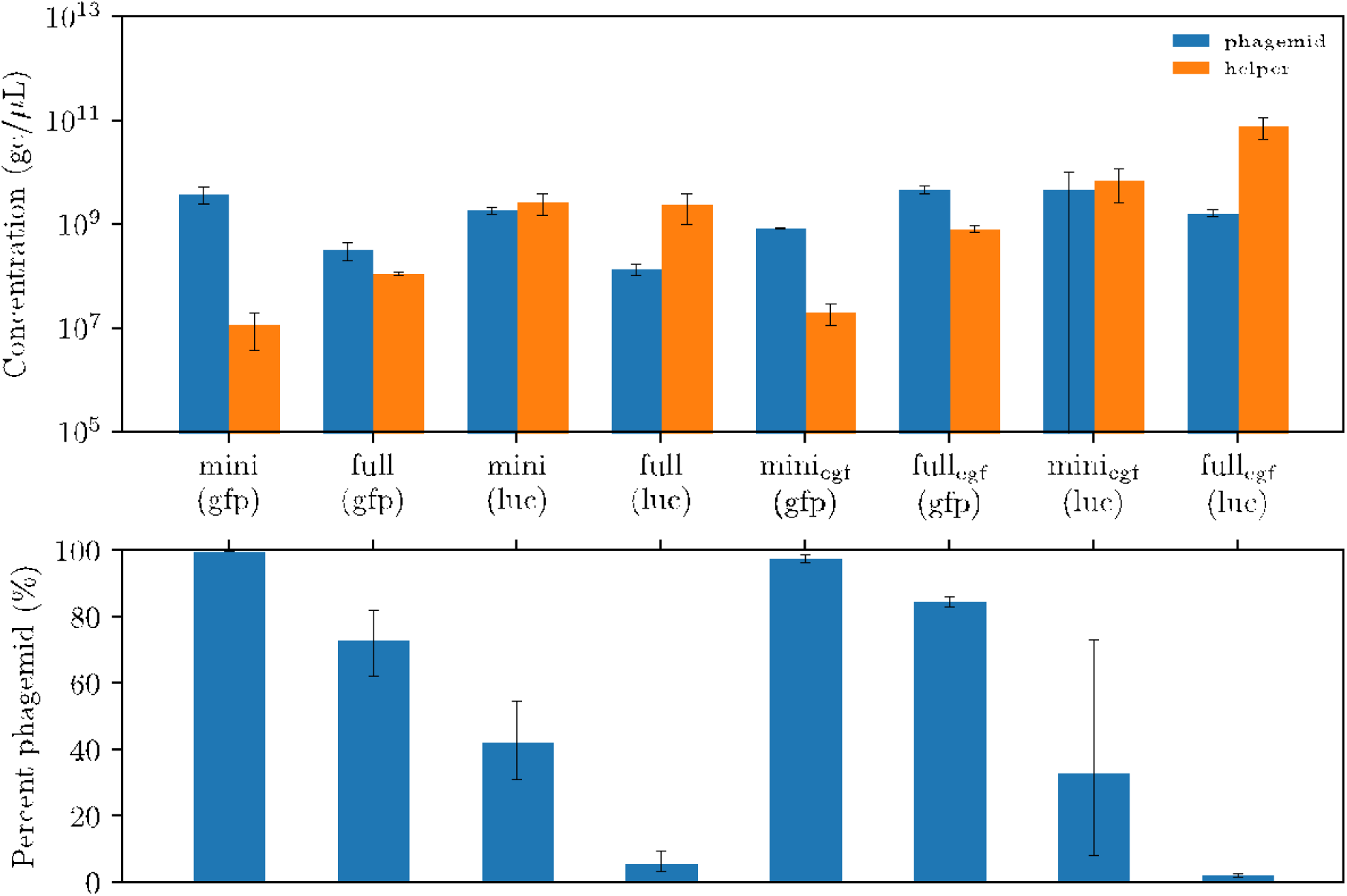
Composition of EGF-displaying phage lysates. At the top, the concentration (gc/μL) of each phage species (target phagemid or helper) in each lysate is presented. Below, the phagemid fraction of each phage species is shown as a percentage of the total phage population. Phagemids on the left were rescued by helper phage M13KO7 (EGF^−^), while phagemids on the right were rescued by helper phage M13SW7-EGF (EGF^+^). Error bars represent SD, *n*=3.

M13 phage particles also did not adversely impact mammalian cell viability. This was regardless of EGF display or complexation with a commercial cationic polymer (TurboFect) both at 24 h and 96 h after transfection (Figure 3A). Phage particles did not reduce cell viability to any significant degree in comparison with the delivery of purified DNA. Purified plasmid dsDNA and phagemid ssDNA were transfected over a period of 96 h in the HeLa cell line (Figure 3B), using the empty backbone vector pM13ori2 as a negative control, and pGL3-CMV (the source of the *cmv-luc* cassette) as a positive control. Luciferase activity peaked approximately 24 to 48 h post-transfection and no significant difference was observed between any of the three dsDNA plasmid vectors. Overall, gene expression, as measured by luciferase activity, was approximately 100-fold lower for purified ssDNA compared to each plasmid counterpart with expression peaking approximately 24 h later, at 72 h. Notably, transfection of purified mini-(*luc*) ssDNA was correlated with increased luciferase activity over purified full-(*luc*) ssDNA. Similar trends were observed when transfecting intact phage particles. Display of EGF dramatically increased gene expression for both mini and full phagemid vectors, which underlines the necessity of receptor-mediated cell internalization for phage-mediated gene transfer.

**Figure 3.**
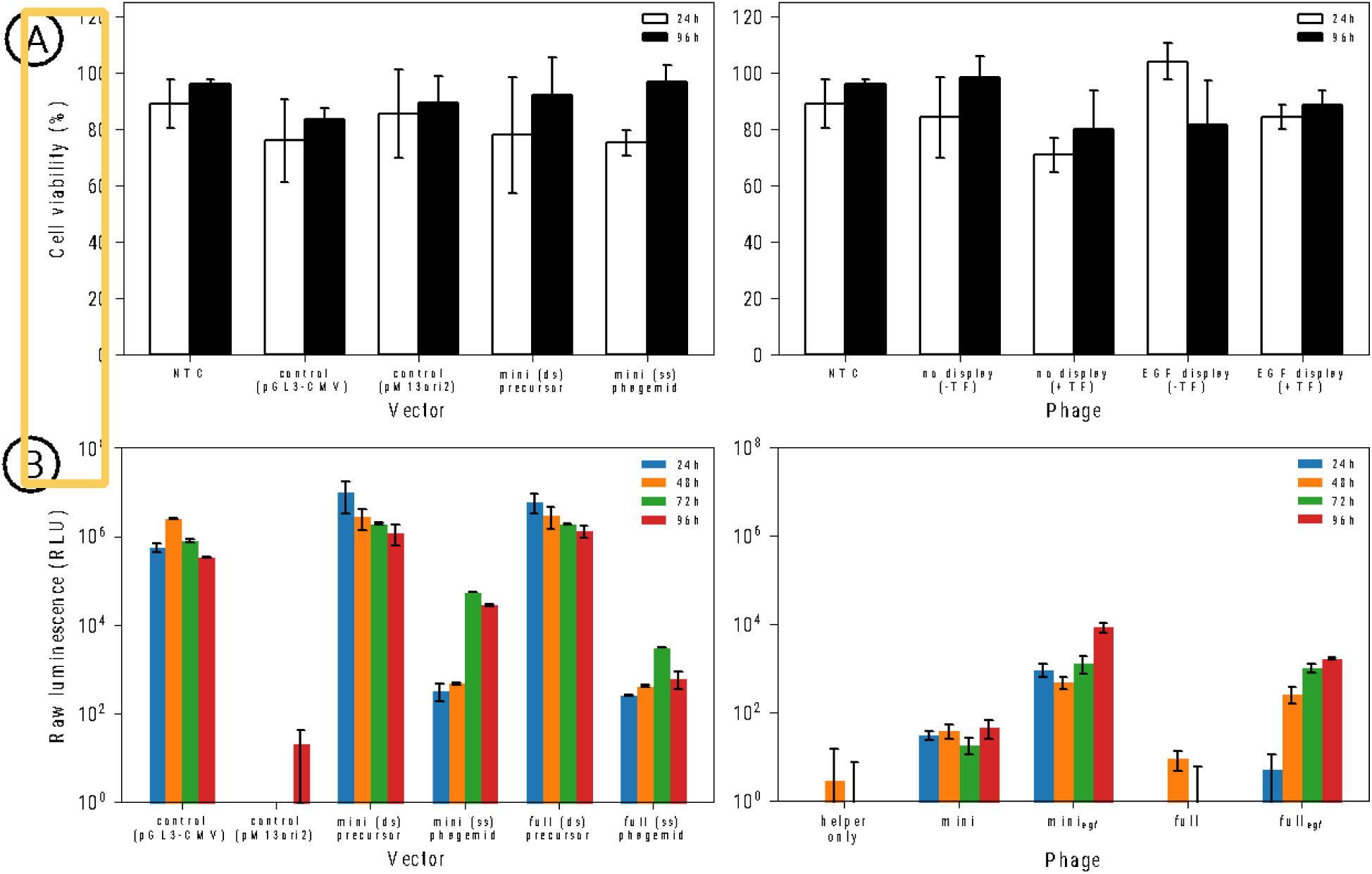
Cell viability is unaffected while transgene expression increases after vector administration. Purified DNA (left) and phage (right) were administered to HeLa cells. A) Cell viability was assessed through the MTT assay 24 and 96 h post-transfection, while B) gene expression was assessed through a luciferase assay. Raw luminescence was reported 24 h, 48 h, 72 h, and 96 h after transfection. The plasmid precursor pM13ori2.cmvluc (mini precursor) and the corresponding purified ssDNA of mini-(luc) (miniphagemid) were transfected alongside the source plasmid pGL3-CMV and the empty vector pM13ori2 as controls. Helper phage M13KO7 (no display) and M13SW7-EGF (EGF display) were transfected alone or with TurboFect. Purified dsDNA was transfected at 1 μg/mL, ssDNA at 2 μg/mL, phage particles at 5 × 10^7^ virions/mL. NTC: no treatment control, TF: TurboFect. Error bars represent SD, *n*=3.

To further characterize the necessity of a targeting ligand, HeLa cells were treated with EGF^+^ or EGF^−^ phage for 1 h, 6 h, 24 h, 48 h, 72 h, and 96 h. The use of EGF as a cell-targeting ligand has previously been shown to improve gene transfer by filamentous phage both *in vitro* and *in vivo*.^10,11,24,25^ Upon ligand binding, internalized EGFR-EGF complexes are routed to lysosomes for degradation; therefore, EGFR-bound phage may be prone to accumulate within juxtanuclear lysosomes, which positions them perfectly for subsequent escape and nuclear transport. Alexa Fluor-tagged M13SW7-EGF abundantly localized to HeLa cells within an hour of administration in a ligand-dependent manner, while M13KO7 did not associate with HeLa cells in any appreciable levels (Figure 4). After 6 h, EGF-displaying phage surrounded the nucleus, indicating successful cell uptake and cytoplasmic translocation. It is known that EGF-EGFR complexes can internalize within 15–20 min^26^, while receptor-bound phages can internalize as early as 10–60 min after administration.^27–30^ Our results appear consistent with these findings.

**Figure 4.**
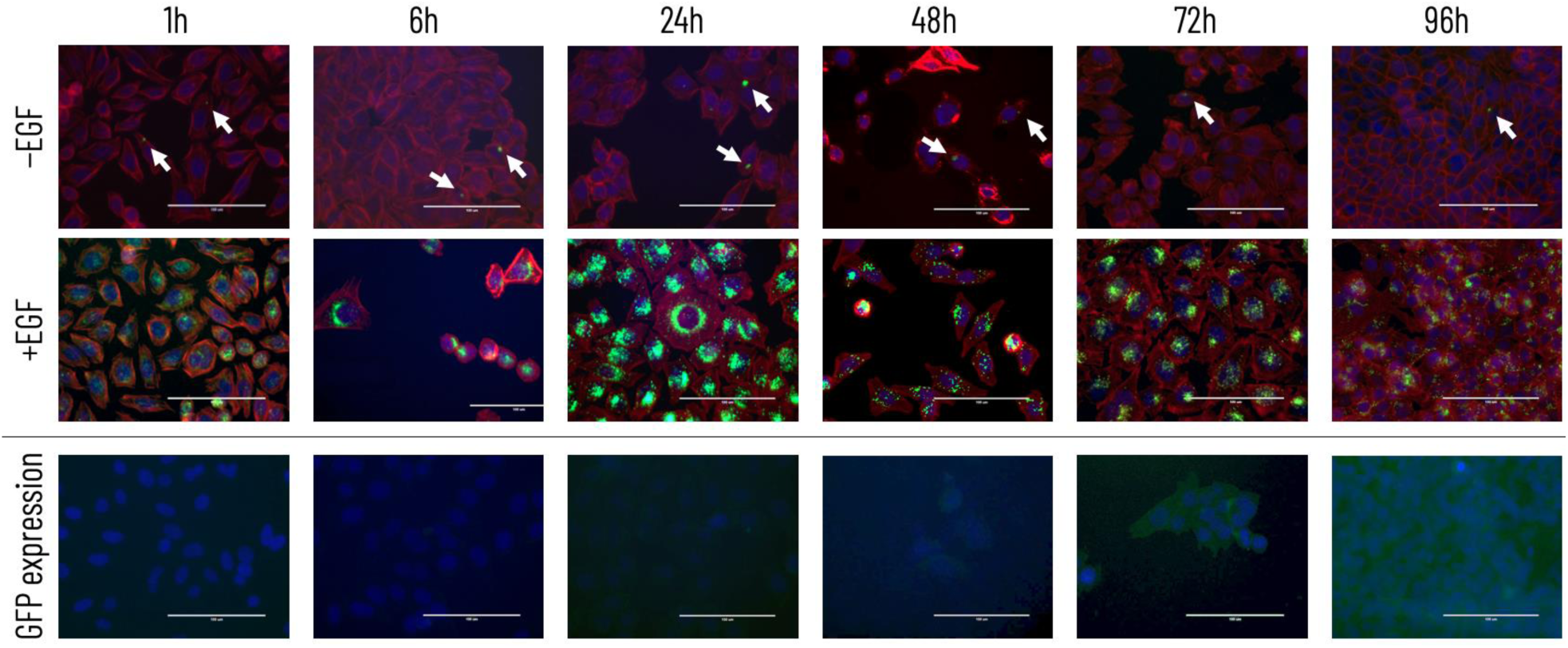
Localization of phage over time in HeLa. M13KO7 (−EGF) and M13SW7-EGF (+EGF) were tagged with Alexa-Fluor 488 (green; top two rows) and applied to HeLa cells, then visualized between 1 to 96 h. Cell nuclei were stained with DAPI (blue) and the cytoskeleton stained with rhodium phalloidin (red). In the bottom row, GFP expression (green) was visualized after administration of mini_egf_-(*gfp*) phage. Nuclei were stained with DAPI.

### Expression efficiency of miniphagemid-mediated gene delivery

The capacity of M13SW7 mini and full phagemids for gene transfer was then compared across four cell lines from different tissues known to moderately express EGFR:^32–37^ HeLa, HT-29, MRC5, and A549, as well as in an EGFR^−^ cell line, HEK293T.^32,38,39^ Firstly, it can be noted that miniphagemids confer increased gene expression compared to full phagemids as an approximately threefold increase was observed for most of the cell lines tested (Table 1). In EGFR^+^ cells, the biggest factor contributing to increased gene expression was the display of the cell-specific ligand (Table 2; Figure 5). This is expected, as the phage must first be taken up through receptor-ligand interactions before any benefits from improved cytoplasmic trafficking can manifest. Although the display of the receptor-targeting ligand was necessary for gene expression, it alone was insufficient for high levels of transgene expression. The rate of phage internalization clearly relies on a number of different factors relating to both the phage used and the targeted tissue type.^27^ Fold differences in luciferase activity between EGF-displaying and non-displaying counterparts showed a dramatic increase in gene expression from the display of EGF (Table 2; Figure 5), indicating a requirement for receptor-mediated vector internalization when the receptor was present. Gene expression was maximally 700-fold greater in HeLa, and over 100-fold greater in HT-29 when the miniphagemid displayed EGF. Overall, our results are generally consistent with previous reports of EGF-mediated improved phage internalization and subsequent gene transfer.^24,25,45^

**Figure 5.**
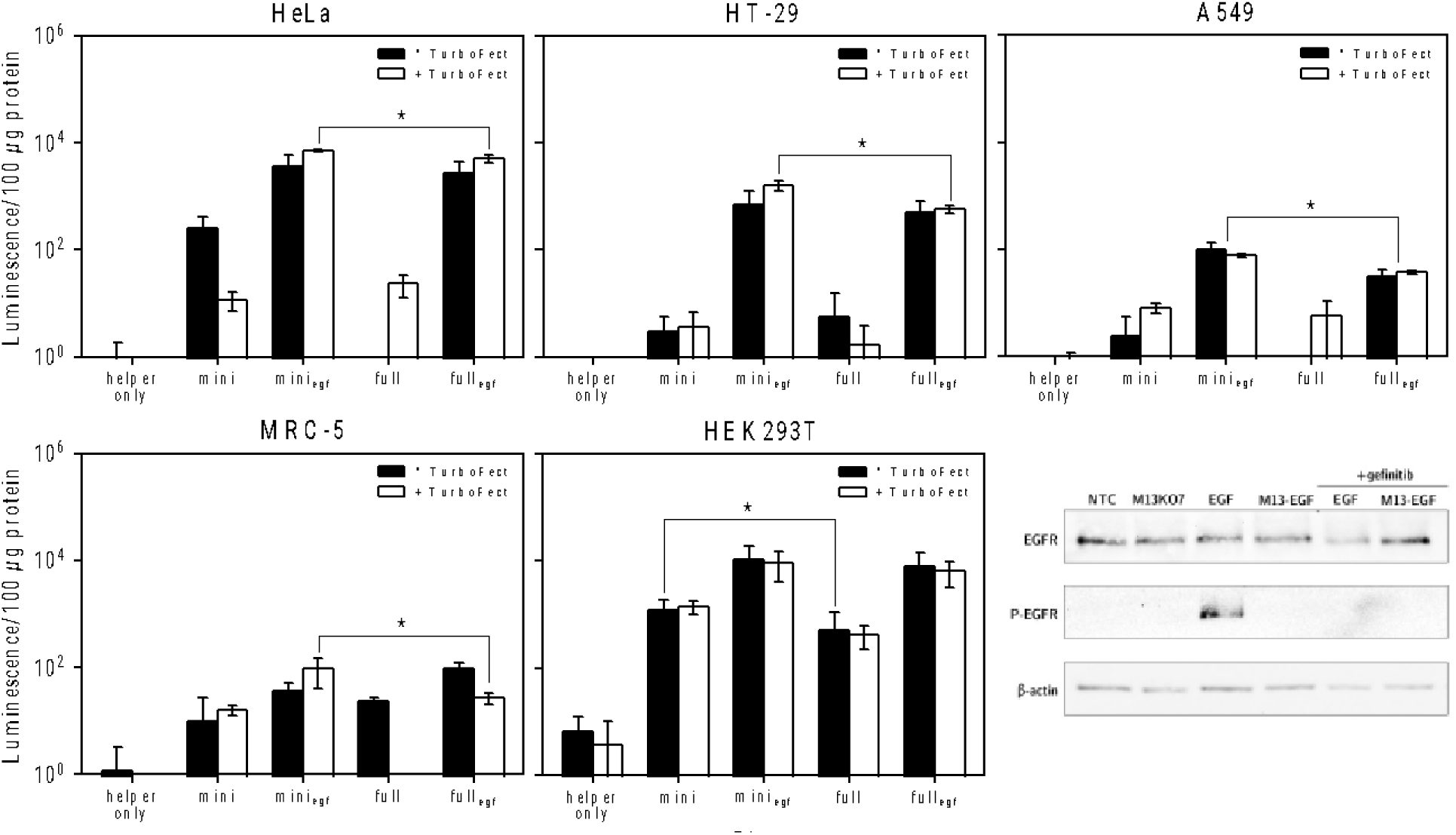
Miniphagemids confer improved transgene expression across EGFR^+^ cell lines. Transgene expression was assessed by luciferase assay 96 h after administration of M13SW7 mini or full phagemids encoding *cmv-luc*. Error bars represent SD, *n*=3. The ∗ above the bars indicates a difference at significance level *p* < 0.05. EGFR phosphorylation was characterized in HeLa cell extracts (Western blot, bottom-right), after administration of M13KO7 (no display), purified recombinant EGF (100 ng/μL) or M13SW7-EGF (EGF display) for 5 min prior to analysis. Additionally, cells were also pre-treated with the EGFR inhibitor, gefitinib (10 μM), prior to treatment with EGF or M13SW7-EGF. Cell lysates were probed for presence of EGFR (EGFR) or phosphorylated EGFR (P-EGFR). β-actin was used as the loading control.

**Table 1.** Fold difference in luciferase expression between M13SW7 mini and full phagemids, *n*=3.

| Cell line | Fold difference in gene expression (mini/full) |  |  |  |
| --- | --- | --- | --- | --- |
|  | EGF<sup>-</sup> |  | EGF<sup>+</sup> |  |
|  | -TurboFect | +TurboFect | -TurboFect | +TurboFect |
| **HEK293T** | 5.22 ± 5.63 | 4.21 ± 3.79 | 2.03 ± 2.59 | 1.78 ± 1.79 |
| **HeLa** | ∞ | 0.51 ± 0.12 | 1.98 ± 1.92 | 1.48 ± 0.37 |
| **HT-29** | ∞ | 7.44 ± 7.63 | 1.74 ± 1.40 | 2.81 ± 0.67 |
| **MRC-5** | 0.52 ± 1.04 | ∞ | 0.43 ± 0.33 | 3.51 ± 1.73 |
| **A549** | ∞ | ∞ | 3.20 ± 0.05 | 2.08 ± 0.18 |

∞: Luminescence below threshold for full phagemid.

**Table 2.** Fold difference in luciferase expression between M13SW7 EGF-displaying and non-displaying phage, *n*=3.

| Cell line | Fold difference in gene expression (EGF<sup>+</sup>/EGF<sup>-</sup>) |  |  |  |
| --- | --- | --- | --- | --- |
|  | mini |  | full |  |
|  | -TurboFect | +TurboFect | -TurboFect | +TurboFect |
| **HEK293T** | 11.73 ± 11.07 | 7.17 ± 4.24 | 43.94 ± 59.44 | 17.65 ± 10.54 |
| **HeLa** | 25.59 ± 33.01 | 704.94 ± 351.36 | ∞ | 261.37 ± 182.45 |
| **HT-29** | ∞ | ∞ | ∞ | ∞ |
| **MRC-5** | ∞ | 5.73 ± 2.79 | 4.14 ± 0.47 | ∞ |
| **A549** | ∞ | 9.96 ± 3.34 | ∞ | ∞ |

∞: Luminescence below threshold for EGF<sup>-</sup> phagemid.

Considering miniphagemids conferred higher gene expression, we next examined M13SW8 and M13SW8-EGF miniphagemids, packaged with mini-(*luc*) ssDNA, for gene expression efficiency in HEK293T(EGFR^-^) and HeLa cells (EGFR^+^). We previously demonstrated that the loss of the PS in the helper phage genome of M13SW8 reduced helper self-packaging, thereby increasing the proportion of miniphagemids carrying the transgene cassette of interest within the miniphagemid lysate.^18^ Lysates of both M13SW8 and M13SW8-EGF miniphagemids showed negligible levels of contaminating phagemid particles, indicating little to no contaminating helper phage or full precursor DNA (Supplementary Figure 2; Supplementary Table 6). An increase in gene expression was observed when treated with EGF-displaying M13SW8 miniphagemids in comparison to non-EGF-displaying M13SW8 miniphagemids (Figure 6) at 96 h, as observed for M13SW7 (Figure 5). Fold differences in gene expression between M13SW8 and M13SW7 showed a range of approximately 40 to 2000-fold increase in HeLa and HEK293T cells (Supplementary Table 7) for EGF-displaying and non-EGF-displaying miniphagemids. Overall, these results demonstrate increased levels of gene delivery when using M13SW8 as compared to M13SW7, suggesting that improvements in the packaging efficiency of DNA minivectors promotes increased gene delivery and expression.

**Figure 6.**
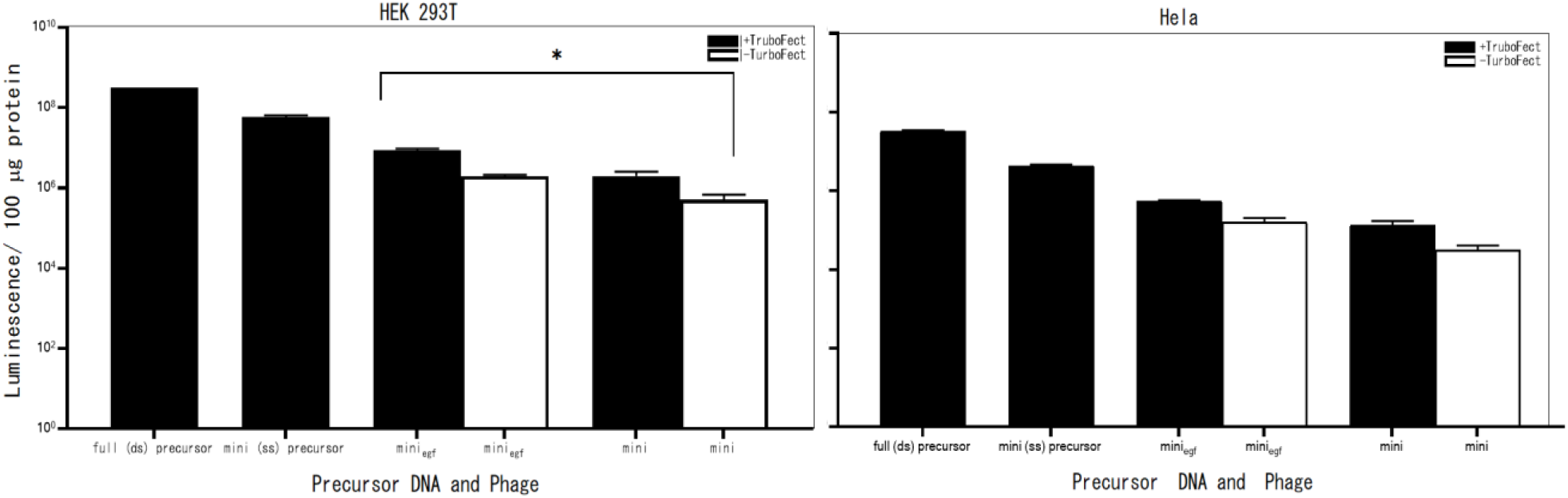
M13SW8-packaged miniphagemids confer improved transgene expression across EGFR^+^ cell lines. Transgene expression was assessed by luciferase assay 96 h after treatment of M13SW8 miniphagemids encoding *cmv-luc* to HEK293T and HeLa cells. Error bars represent SD, n=3. The ∗ above the bars indicates a difference at significance level *p* < 0.05, only recorded for comparison between the miniphagemid samples (with and without transfection reagent).

### Filamentous phage uptake is independent of EGFR phosphorylation

Despite the use of EGFR as an internalization target in multiple studies, activation of EGFR-mediated signal transduction by EGF-displaying phage has not been deeply investigated. Canonically, ligand-EGFR interactions lead to receptor dimerization and receptor autophosphorylation prior to internalization of the entire receptor-ligand complex via clathrin-coated endocytosis.^47,48^ Activation of the EGF receptor stimulates signal transduction pathways involved in cell proliferation. Phosphorylation of several intracellular tyrosine residues of EGFR mediates downstream signal transduction. After treatment of HeLa cells with purified recombinant EGF, phosphorylated EGFR was detectable but not if cells were pre-treated with gefitinib, an EGFR inhibitor (Figure 5). In contrast, no phosphorylation was detected after treatment with EGF-displaying phage. The M13 phage particle itself appears sufficient to prohibit EGFR ligand-activated autophosphorylation, without preventing ligand-activated internalization.

## Discussion

### Display of EGF does not impact phage packaging or infectivity

As pIII is a minor coat protein and does not participate in phagemid replication or ssDNA sequestration, it was not expected to impact miniphagemid production. Indeed, quantification of the phage lysates showed that the fusion did not inhibit helper phage rescue of phagemid from either a split or wild-type origin (Figure 2; Supplementary Table 5). Since the display fusion was encoded on the helper phage, other fusion peptides could be incorporated without modifying the phagemid vector itself. For chimeric display on other coat proteins (ex: pVIII), it may be possible to incorporate a fusion peptide on the backbone of the phagemid vector such that it would not be assembled into progeny viral particles.

### Display of EGF does determine phage cellular uptake

The strong perinuclear accumulation of tagged phage particles suggests they have not yet escaped the endosomal compartment at 6 h, as escaped phage particles are expected to localize more diffusely throughout the cytoplasm.^30^ Still, expression of phage-encoded transgene cassettes was detectable 48-72 h post-treatment, which was on par with the delivery of purified ssDNA. Either the additional requirement of DNA separation from the filamentous phage coat does not significantly delay phage-mediated gene transfer or potential improvements from phage-mediated cell uptake and intracellular navigation are sufficient to offset delays. Indeed, the low luminal pH of juxtanuclear lysosomes can contribute to phage coat shedding,^31^ possibly improving the bioavailability of phage-encapsulated DNA.

### The single-stranded miniphagemid improves gene transfer

Removal of the bacterial backbone conferred an increase in gene expression as compared to the full phagemid with an intact bacterial backbone for M13SW7 phagemids (Table 1; Figure 5). This increase in gene expression was statistically significant in combination with the display of EGF and when complexed with TurboFect in all four EGFR^+^ cell lines (*p* < 0.05). In HEK293T, phagemid miniaturization correlated with increased luciferase activity when complexed with TurboFect (*p* < 0.05), but there was no statistically significant difference between phagemids with or without ligand display. However, in HEK293T cells treated with M13SW8 miniphagemids, statistical significance (p < 0.05) was observed for miniphagemids displaying EGF and complexed with TurboFect, compared to non-displaying miniphagemids without complexing with TurboFect (Figure 6). Phage uptake may have occurred independently of the targeting ligand, indicating a potential for nonspecific tissue uptake. This contradicts the observed inability of EGF^−^ phage to transfect EGFR^+^ cell lines. It may be that HEK293T cells express a cell surface receptor with intrinsic affinity for M13; however, studies of M13 in mice have not shown strong preferential accumulation in the kidney.^40^ Filamentous phage particles have been previously reported to enter via caveolae-mediated endocytosis^41,42^, while larger phage particles enter via phagocytosis and micropinocytosis.^43^ In the absence of a target receptor, filamentous phage uptake in HEK293T is likely clathrin-independent, but the specific endocytic mechanism and how the phage targets the cell remains unclear and requires further investigation.

Intriguingly, phage-mediated gene expression was slightly improved when combined with a cationic polymer. This is consistent with the observations of Donnelly et al., who reported that the cationic polymer PEI improved filamentous phage-mediated gene transfer.^44^ Complexation with a cationic polymer enhanced EGF-dependent gene transfer but did not permit EGF-independent gene transfer, suggesting that its benefits may be realized intracellularly. This is more indicative of a role in facilitating endosomal escape rather than cell entry.

### Internalization of EGF-displaying M13 does not activate EGFR

Larger molecules such as the monoclonal antibody cetuximab^49^ and bacteriophage λ^50^ have been shown to internalize through EGFR binding without triggering the downstream signalling pathway.^51^ Antagonistic receptor binding by cetuximab and λ have both been shown to reduce cell proliferation. However, we and others have not observed a decrease in cell viability over time after administration of EGF^+^ filamentous phage. Intriguingly, the induction of late EGFR-stimulated events has been reported with other EGF-displaying M13: specifically, M13 display of EGF was involved in the activation of c-fos serum response element (SRE)-mediated transcription.^52^ Still, it has been demonstrated elsewhere that these events can occur even in cells with kinase-defective EGFR^53^, suggesting alternative mechanisms to stimulate EGFR-mediated signal transduction separately from the EGF receptor specifically. Downstream cell proliferation pathways could be activated in the absence of cell surface EGFR phosphorylation.^54–56^ While we did not observe phage-mediated EGFR autophosphorylation, this does not rule out endosomal EGFR signalling nor other kinase-independent signal transduction. Further investigations with EGF-displaying phage are warranted.

Overall, both purified and phage-encapsulated miniphagemid DNA were correlated with greater gene expression over their full phagemid counterparts, which we attributed to the reduction in immunostimulatory CpG motifs and improved cytoplasmic diffusion. Another benefit conferred by the smaller size of the phagemid particle may be more efficient internalization upon ligand-receptor binding. Clathrin-mediated endocytosis has been previously shown to accommodate filamentous phage particles up to 900 nm in length^30^, even though clathrin-coated vesicles are typically on the scale of 200 nm^46^. Although long, their flexible rod-like structure enables filamentous phages to be compacted; as such, they are more readily taken up alongside receptor-mediated clathrin or caveolae-mediated endocytosis in contrast to other larger, more globular proteins and phages. This effect is likely enhanced by the shorter nature of the miniphagemid particles since filamentous phage length is determined largely by the length of the encapsulated DNA molecule.

### Deletion of PS in the helper plasmid improves packaging efficiency and gene transfer

We previously demonstrated improved miniphagemid rescue efficiency via deletion of the PS in the helper phage genome^18^, which we additionally confirmed here (Supplementary Figure 2; Supplementary Table 6). Significant packaging of the helper genome is a common issue with helper phages such as M13SW7 and M13KO7. Phage genomes in the resultant lysate adds contaminating, potentially immunogenic bacterial DNA, and reduces the concentration of transgene-encoding molecules per phage lysate. Consequently, the system was optimized by deleting the PS of M13KO7, generating M13SW8, in order to prevent packaging of self-DNA^18^. Miniphagemids produced using M13SW8 would therefore exhibit enhanced delivery and gene expression as a higher percentage of phagemids produced are target miniphagemids. When HEK293T and HeLa cells were transfected with M13SW8 miniphagemids an increase in luciferase expression compared to M13SW7 miniphagemids was observed (Supplementary Table 7), indicating that a self-packaging deficient helper phage resulted in superior lysates for transfections. This is most likely due to an increased amount of target miniphagemids delivered to target cells.

## Summary and conclusions

Our work here underscores the power of phage display as a means to confer tropism for phage-mediated gene transfer in mammalian cells. By incorporating a separate helper phage that encodes the display fusion, miniphagemids displaying any peptide of interest can be propagated very easily. We described here the use of EGF-displaying phage encoding luciferase, but the combination of other cell-targeting ligands and transgene cassettes makes for a powerfully flexible vector platform. The complexation of phage particles with a cationic polymer also further enhances gene transfer; we postulate this effect likely manifests by mediating escape from the endosomal compartment after phages have already internalized. Future developments in bacteriophage gene delivery must address the need for endosomal escape in order to increase intracellular phagemid bioavailability. Importantly, we have shown that removal of the bacterial backbone in miniphagemid vectors improved M13-mediated gene transfer up to threefold across multiple different cell lines, likely by increasing cytoplasmic diffusion towards the nucleus and improving immune evasion through the loss of inflammatory bacterial andphage DNA sequences. Additionally, the deletion of the packaging signal in the helper phage plasmid further improved packaging efficiency, which will be helpful in scaled-up production of this system, in addition to improving M13-mediated gene transfer of up to 2000-fold. Overall, combining results obtained for M13SW7 and M13SW8 phagemids, we have demonstrated improved gene delivery through the combination of cell-targeting ligand display (EGF), helper plasmid self packaging minimization, and vector minimization.

## Data Availability

All relevant data are within the manuscript.

## Acknowledgements

Many thanks to J. Blay, M. Aucoin, and N. Oviedo for their generosity in sharing materials.

## Author Contributions

Conceptualization, S.W., S.J., and R.S.; Methodology, S.W., S.J., and R.S.; Formal Analysis, S.W.; Investigation, S.W., S.J., D.P., R.P.; Visualization: S.W.; Writing – Original Draft Preparation, S.W.; Writing – Review & Editing, S.J, R.S., D.P., R.P.; Supervision, R.S.; Funding Acquisition, R.S.

