## Supplementary Data (Figures 1-2 and Tables 1-7) for "A novel miniaturized filamentous phagemid as a gene delivery vehicle to target mammalian cells"

### Supplementary Information

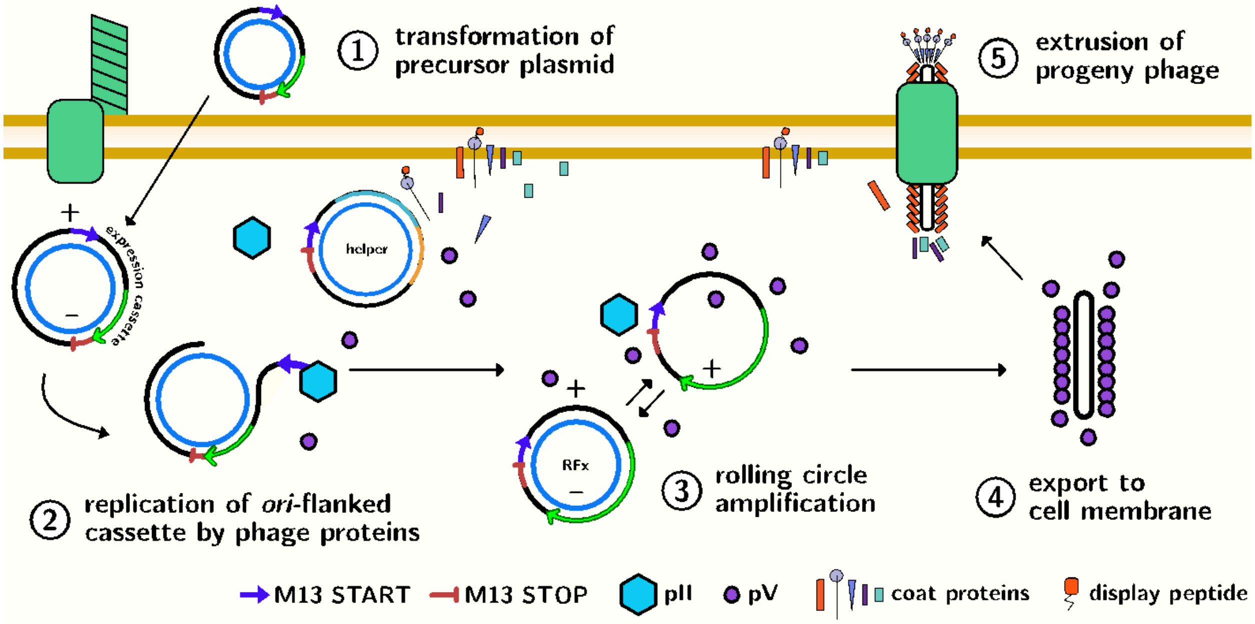


**Figure S1: Production and purification of miniphagemid particles.** 1) The host cell is transformed by the precursor plasmid encoding a region of interest between the separated M13 replication signals. 2) After infection with a helper phage encoding a display peptide, phage protein expression can occur. 3) Rolling circle amplification generates a recombinant replicative factor (RFx) from the plasmid that reconstitutes the f1 *ori* and loses the plasmid backbone. 4) The phage ssDNA binding proteins pV sequester single-stranded minivector, preventing further amplification, and shift the replication cycle towards assembly. 5) Assembly proteins extrude progeny phage particles encapsulating the minivector that homogeneously display a peptide of interest (Wong et al., 2023).

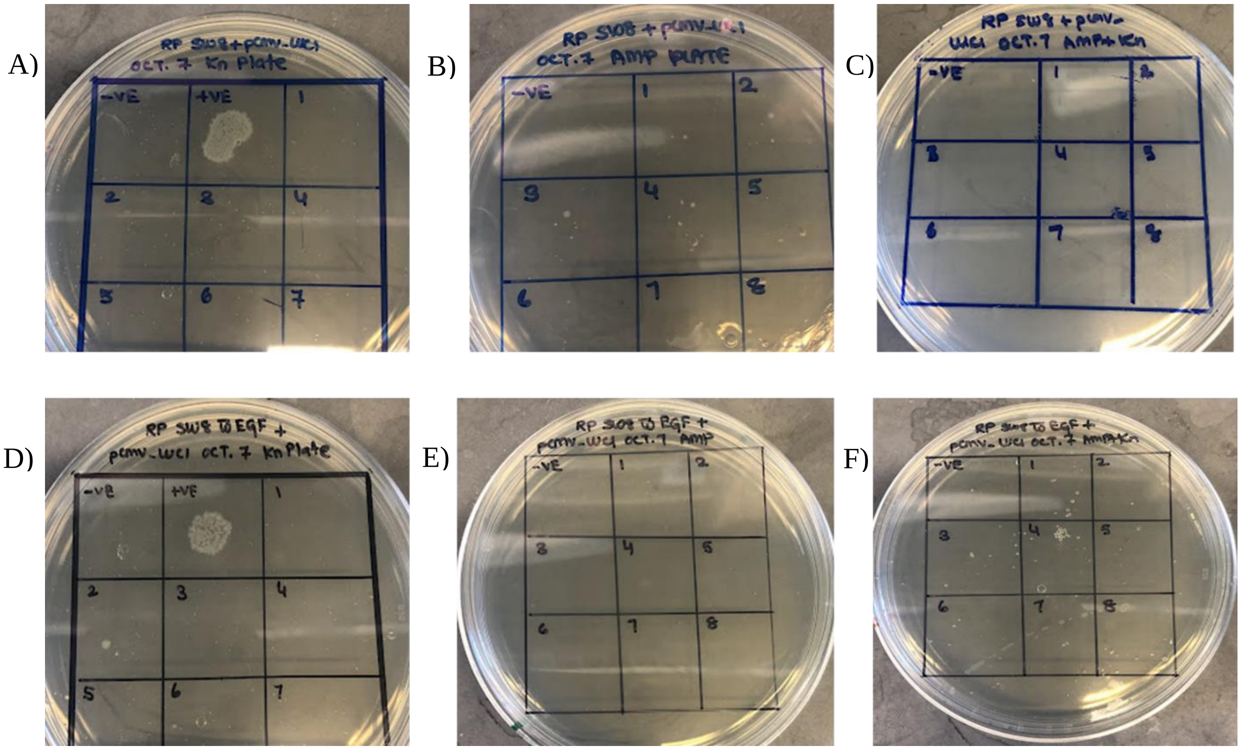


**Figure S2: Efficiency of plating spot plates.** M13SW8-mini-(luc) spot plates on a) LB + kanamycin; b) LB + ampicillin; c) LB + ampicillin + kanamycin. M13SW8-mini$_{egf}$-(luc) spot plates on d) LB + kanamycin; e) LB + ampicillin; f) LB + ampicillin + kanamycin. 10 uL of phage dilutions in TN buffer were plated starting at a titre of $10^{-2}$ to $10^{-8}$ (labelled on plates as 1-7). The positive controls were 10 uL of plasmid containing ampicillin or kanamycin resistance genes diluted in TN buffer. The negative control was 10 uL of TN buffer.

**Table S1:** Strains used in this study

| Strain | Genotype/description | Source |
|---|---|---|
| **Bacterial strains** | | |
| JM109 | F′ *traD36 proAB+ lacIq lacZΔM15/ Δ(lac-proAB) endA1 glnV44 thi-1 e14− recA1 gyrA96 relA1 hsdR17* | New England BioLabs |
| **Mammalian cell lines** | | |
| HEK-293T | Embryonic kidney, epithelial | Gift, Dr. M. Aucoin |
| HeLa | Uterus, cervix adenocarcinoma | Gift, Serenity Bioworks |
| MRC-5 | Lung, fibroblast | ATCC CCL-171 |
| HT-29 | Colon, epithelial adenocarcinoma | Gift, Dr. J. Blay |
| A549 | Lung, epithelial carcinoma | ATCC CCL-185 |

**Table S2:** Plasmids used in this study

| Plasmid | Genotype | Source |
|---|---|---|
| pGL2-SS-CMV-GFP-SS | pGL2-Promoter, *cmv-gfp* replaces SV40-*luc*, AmpR | Gift, Mediphage Bioceuticals |
| pGL3-CMV | pGL3-Basic, *cmv* inserted in BlgII-HindIII, AmpR | Gift, Dr. N. Oviedo [57] |
| pBluescript II KS+ | Wild-type f1 *ori*, pUC *ori*, AmpR | |
| M13SW8 | M13KO7, Packaging signal removed, $Kn^R$ | Our previous study[18] |
| M13SW8-EGF | M13SW8, EGF display on pIII, $Kn^R$ | Our previous study[18] |
| M13SW7 | M13KO7, gIII from M13KE | This study |
| M13SW7-EFF | M13SW7, *egf* inserted in KpnI-EagI | This study |
| pSW9 | pBluescript II KS+, *cmv-gfp* inserted in KpnI, AmpR | Our previous study[18] |
| pSW10 | pBluescript II KS+, *cmv-luc* inserted in KpnI, AmpR | Our previous study[18] |
| pM13ori2 | pUC57, M13-START and M13-STOP in *lacZa* | Our previous study[18] |
| pM13ori2.cmvgfp | pM13ori2, *cmv-gfp* from pGL2-SS-CMV-GFP-SS inserted in EcoRI-PacI | Our previous study[18] |
| pM13ori2.cmvluc | pM13ori2, pM13ori2, *cmv-luc* from pGL3-CMV inserted in EcoRI-KpnI | Our previous study[18] |

**Table S3:** Phages used in this study

| Phage | Genotype | Source |
|---|---|---|
| M13KO7 | Tn903 (p15a *ori*, KanR) | New England BioLabs |
| M13KE | lacZα, KpnI & EagI in gIII | New England BioLabs |
| M13SW7 | M13KO7, gIII from M13KE | This study |
| M13SW7-EGF | M13SW7, *egf* inserted in KpnI-EagI | This study |
| M13SW7-full-(gfp) | from precursor pSW9, AmpR | This study |
| M13SW7-full$_{egf}$-(gfp) | from precursor pSW9, AmpR | This study |
| M13SW7-full-(luc) | from precursor pSW10, AmpR | This study |
| M13SW7-full$_{egf}$-(luc) | from precursor pSW10, AmpR | This study |
| M13SW7-mini-(gfp) | from precursor pM13ori2.cmvgfp, AmpR | This study |
| M13SW7-mini$_{egf}$-(gfp) | from precursor pM13ori2.cmvgfp, AmpR | This study |
| M13SW7-mini-(luc) | from precursor pM13ori2.cmvluc, AmpR | This study |
| M13SW7-mini$_{egf}$-(luc) | from precursor pM13ori2.cmvluc, AmpR | This study |
| M13SW8-mini-(luc) | from precursor pM13ori2.cmvluc, AmpR | This study |

**Table S4:** Primers for M13SW7 and M13SW7-EGF phage construction

| Primer | Amplicon | Sequence (5′ − 3′)[1] |
|---|---|---|
| gIII-F | gIII | TTTTTTTGGAGATTTTCAACGTGAAAAAATTATTATTCGCA ATTCC |
| gIII-R | gIII | CCCAAAAGAACTGGCATGATTTAAGACTCCTTATTACGCAG TATG |
| M13KO7-F | M13KO7 | GTTGAAAATCTCCAAAAAAAAAGGC |
| M13KO7-R | M13KO7 | TCATGCCAGTTCTTTTGGG |
| KpnI-egf-F | *egf* | *GGTACCTTTCTATTCT*CACTCTAATAGTGACTCTGAATGTC CC[2] |
| EagI-egf-R | *egf* | CGGCCGA*AGAACCACCACC*GCGCAGTTCCCACCAC[3] |

[1]Underlined nucleotides indicate primer overhang. [2]Italicized nucleotides indicate gIII leader sequence. [3]Italicized nucleotides indicate GGGS linker.

**Table S5:** Phage titres

| Phage | Titre (x $10^{13}$ PFU/mL)[1] |
|---|---|
| M13 | 0.13 ± 0.05 |
| M13KO7 | 1.90 ± 0.10 |
| M13KE | 0.12 ± 0.02 |
| M13SW7 | 3.43 ± 1.50 |
| M13SW7-EGF | 2.75 ± 0.90 |

**Table S6:**Efficiency of plating (EOP) for M13SW8 and M13SW8-EGF miniphagemids

| Miniphagemid | Titre (PFU/mL) | Ampicillin EOP (%) | Kanamycin EOP (%) | Ampicillin + Kanamycin EOP (%) |
|---|---|---|---|---|
| M13SW8-mini-(luc) | 1 x $10^{13}$ | 2 x $10^{-8}$ | <1 x $10^{-8}$ | <1 x $10^{-8}$ |
| M13SW8-mini$_{egf}$-(luc) | 1 x $10^{12}$ | 1 x $10^{-7}$ | <1 x $10^{-7}$ | <1 x $10^{-7}$ |

Equation for efficiency of plating: (contaminated phagemid (helper phage + full precursor) titre/total phagemid titre) x 100

**Table S7:** Fold differences in gene expression between M13SW8 and M13SW7 miniphagemids, n= 3

| Cell line | Fold difference in gene expression(M13SW8/M13SW7) | | | |
|---|---|---|---|---|
| | EGF+ | | EGF- | |
| | -TurboFect | +Turbofect | -TurboFect | +TurboFect |
| HEK293T | 270.08 | 570.88 | 788.11 | 2131.78 |
| HeLa | 40.08 | 79.03 | 4742.63 | 1939.45 |
